# SARS-CoV-2 mutations acquired in mink reduce antibody-mediated neutralization

**DOI:** 10.1101/2021.02.12.430998

**Authors:** Markus Hoffmann, Lu Zhang, Nadine Krüger, Luise Graichen, Hannah Kleine-Weber, Heike Hofmann-Winkler, Amy Kempf, Stefan Nessler, Joachim Riggert, Martin Sebastian Winkler, Sebastian Schulz, Hans-Martin Jäck, Stefan Pöhlmann

## Abstract

Transmission of SARS-CoV-2 from humans to farmed mink was observed in Europe and the US. In the infected animals viral variants arose that harbored mutations in the spike (S) protein, the target of neutralizing antibodies, and these variants were transmitted back to humans. This raised concerns that mink might become a constant source of human infection with SARS-CoV-2 variants associated with an increased threat to human health and resulted in mass culling of mink. Here, we report that mutations frequently found in the S proteins of SARS-CoV-2 from mink were mostly compatible with efficient entry into human cells and its inhibition by soluble ACE2. In contrast, mutation Y453F reduced neutralization by an antibody with emergency use authorization for COVID-19 therapy and by sera/plasma from COVID-19 patients. These results suggest that antibody responses induced upon infection or certain antibodies used for treatment might offer insufficient protection against SARS-CoV-2 variants from mink.

## INTRODUCTION

The pandemic spread of Severe Acute Respiratory Syndrome Coronavirus 2 (SARS-CoV-2) and the associated disease coronavirus disease 2019 (COVID-19) resulted in 105 million diagnosed infections and 2.3 million deaths ((WHO), 2021). The virus has been introduced into the human population in China in the winter season of 2019, and first cases were detected in the city of Wuhan, Hubei province (Zhou et al., 2020). Bats and pangolins harbor viruses closely related to SARS-CoV-2 and are discussed as sources for SARS-CoV-2 (Lam et al., 2020; Xiao et al., 2020; Zhou et al., 2020). However, it is conceivable that other animals contributed to the spillover of the virus from animals to humans, considering that SARS-CoV was transmitted from bats to humans via civet cats and raccoon dogs (Guan et al., 2003; Lau et al., 2005; Li et al., 2005).

The American mink (*Neovison vison*) is farmed in Denmark, the Netherlands and many other countries for its fur. In April 2020, mink in individual farms in the Netherlands developed a respiratory disease and SARS-CoV-2 was detected in the afflicted animals (Molenaar et al., 2020; Oreshkova et al., 2020). Whole-genome sequencing provided evidence that SARS-CoV-2 was initially introduced into mink from humans and that farm workers subsequently acquired the virus from infected animals (Oude Munnink et al., 2020). Further, the data suggested that viruses acquired from infected mink were capable of human-to-human transmission ((Oude Munnink et al., 2020), comments: (Koopmans, 2020; Leste-Lasserre, 2020)). SARS-CoV-2 infection of farmed mink and transmission of the virus from infected animals to humans was subsequently also detected in Denmark and led to the culling of 17 million animals. Finally, apart from the Netherlands and Denmark, also other countries reported SARS-CoV-2 infections of farmed and free-ranging mink, including several European countries (ProMed-mail, 2020a, b, d, e, f) (Fig. 1A), Canada (ProMed-mail, 2020g) and the USA (ProMed-mail, 2020c, h).

**Figure 1.**
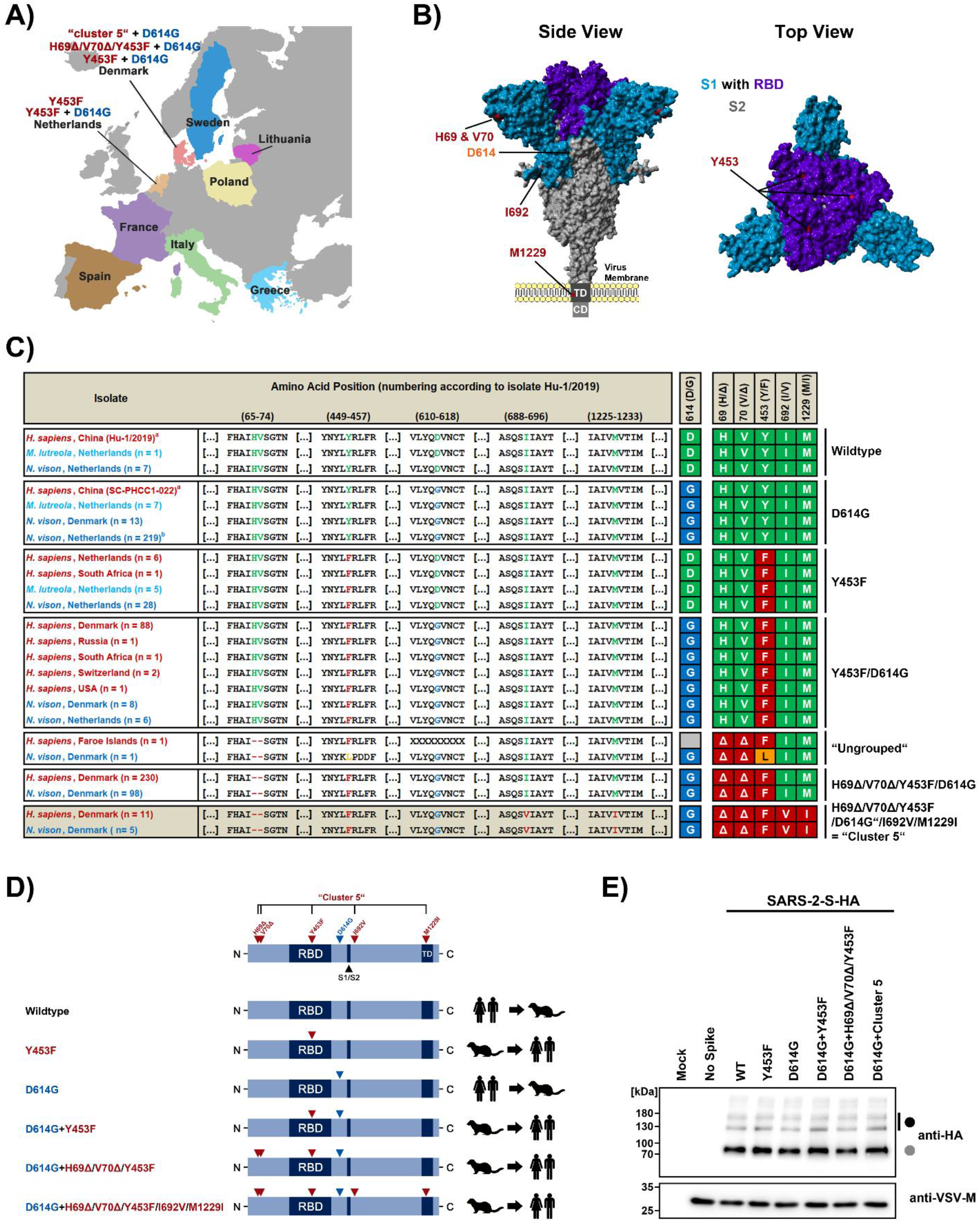
Mink-specific spike protein variants are robustly expressed, proteolytically processed and incorporated into viral particles. (A) European countries that have reported SARS-CoV-2 infection in mink. The mink-specific spike (S) protein mutations under study are highlighted. (B) Summary of mink-specific S protein mutations found in human and mink SARS-CoV-2 isolates. Sequences were retrieved from the GISAID (global initiative on sharing all influenza data) database. Legend: a = reference sequences, b = 36/219 sequences carry additional L452M mutation; Abbreviations: *H. sapiens* = *Homo sapiens* (Human), *N. vison* = *Neovison vison* (American Mink), *M. lutreola* = *Mustela lutreola* (European Mink). (C) Location of the mink-specific S protein mutations in the context of the 3-dimensional structure of the S protein. (D) Schematic illustration of the S protein variants under study and their transmission history. Abbreviations: RBD = receptor binding domain, S1/S2 = border between the S1 and S2 subunits, TD = transmembrane domain. (E) Rhabdoviral pseudotypes bearing the indicated S protein variants (equipped with a C-terminal HA-epitope tag) or no viral glycoprotein were subjected to SDS-PAGE under reducing conditions and immunoblot in order to investigate S protein processing and particle incorporation. Detection of vesicular stomatitis virus matrix protein (VSV-M) served as loading control. Black and grey circles indicate bands for unprocessed and processed (cleavage at S1/S2 site) S proteins, respectively. Similar results were obtained in four separate experiments.

The SARS-CoV-2 spike (S) protein is incorporated into the viral envelope and facilitates viral entry into host cells. For this, the S protein binds to the cellular receptor angiotensin-converting enzyme 2 (ACE2) via its receptor-binding domain (RBD) and employs the cellular serine protease TMPRSS2 for S protein priming (Hoffmann et al., 2020; Zhou et al., 2020). The S protein of SARS-CoV-2 from farmed mink in Denmark and the Netherlands harbors different combinations of mutations relative to SARS-CoV-2 circulating in humans (Oude Munnink et al., 2020) (Fig. 1B and C): A deletion of H69 (H69Δ) and V70 (V70Δ) in the S protein N-terminus and amino acid exchanges Y453F in the RBD, I692V located downstream of the furin motif, S1147L in the S2 subunit and M1229I in the transmembrane domain (Fig. 1B and C). Moreover, SARS-CoV-2 containing a combination of five mutations (H69Δ/V70Δ/Y453F/I692V/M1229I) in their S protein have been observed, which gave rise to the designation cluster 5 variant. Here, we investigated whether S proteins harboring Y453F either alone or in conjunction with other mutations showed altered expression, host cell interactions and susceptibility to antibody-mediated neutralization.

## RESULTS

We employed previously described vesicular stomatitis virus-based reporter particles bearing the SARS-CoV-2 S protein to study whether mutations observed in infected mink modulate cell entry and its inhibition (Hoffmann et al., 2020). The S protein from SARS-CoV-2 isolate hCoV-19/Wuhan/Hu-1/2019, which harbors an aspartic acid at amino acid position 614 (D614) (Korber et al., 2020), was used as control and is subsequently referred to as wildtype (WT). Further, an S protein of identical amino acid sequence but harboring a glycine at position 614 (D614G), was used as a reference for S protein variants containing the dominant D614G mutation (Fig. 1D). Finally, S proteins with mutations found in SARS-CoV-2 from mink were analyzed as shown in figure 1D.

Immunoblot analysis of S protein-bearing particles revealed that all mutations were compatible with robust particle incorporation of the S protein and cleavage at the furin motif located at the S1/S2 cleavage site (Fig. 1E). Similarly, all S proteins efficiently utilized human ACE2 upon directed expression in otherwise non-susceptible BHK-21 cells (Fig. 2A). Further, all tested S proteins mediated entry into cell lines commonly used for SARS-CoV-2 research (Fig. 2B), which were also readily transduced by control particles bearing VSV-G (SI Fig. S1). Substitution D614G, which is dominant in SARS-CoV-2 from humans (Korber et al., 2020) and was also found in viruses from mink, increased the efficiency of S protein-driven entry, as expected (Korber et al., 2020; Plante et al., 2020). Combination of D614G with the mink-specific mutation Y453F (mutant D614G+Y453F) or Y453F in conjunction with H69Δ, H70Δ (mutant D614G+H69Δ/H70Δ/Y453F) did not modulate entry efficiency when compared to D614G alone (Fig. 2B). Finally, mutation D614G+cluster 5 reduced entry into several cell lines but was compatible with robust entry into the human intestinal cell line Caco-2 and the lung cell line Calu-3 (Fig. 2B). Thus, mutations detected in the S proteins of SARS-CoV-2 from mink were compatible with robust viral entry into human intestinal and lung cells.

**Figure 2.**
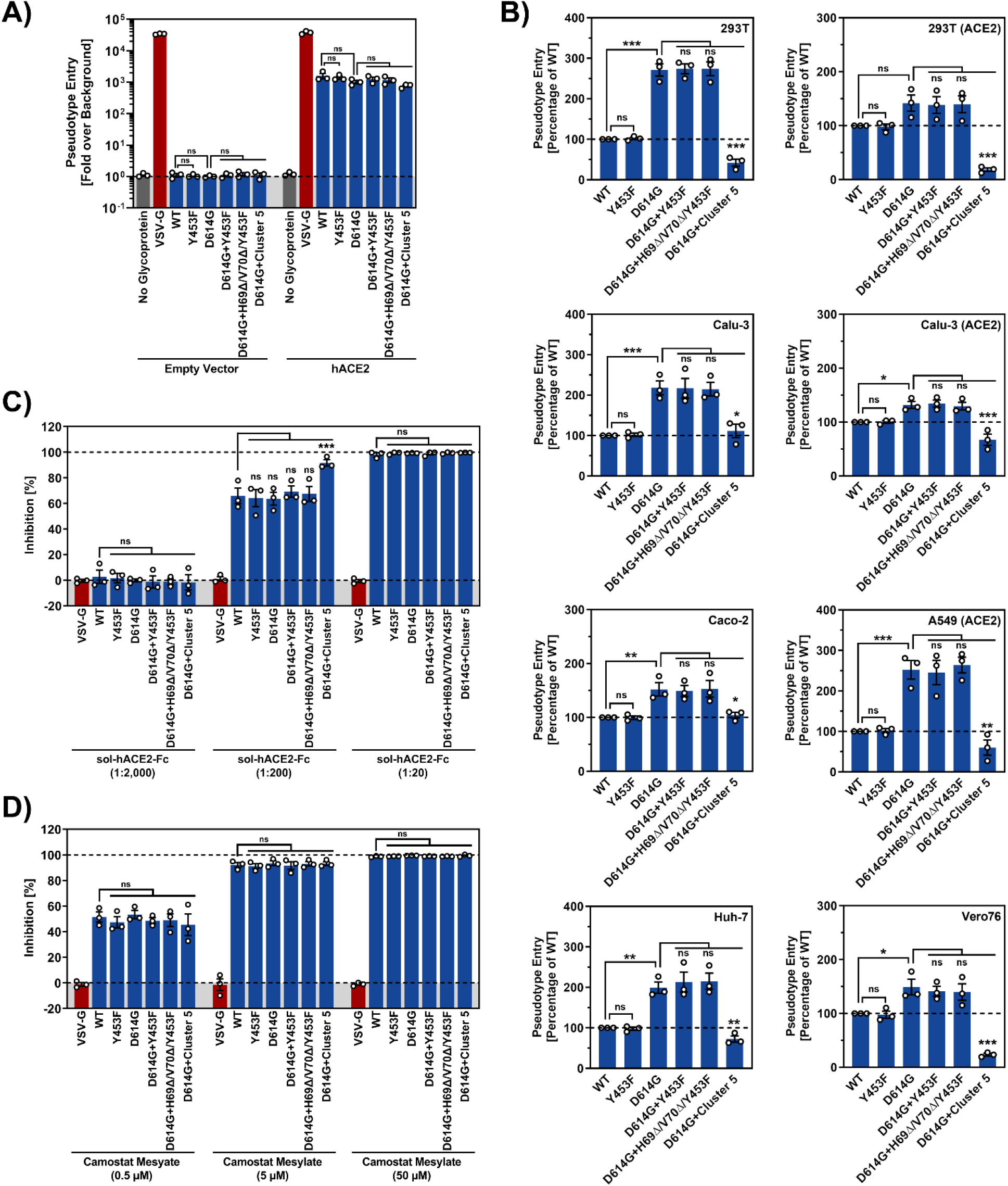
Spike protein variants found in mink enable robust entry into human cells and entry is blocked by soluble ACE2 and the protease inhibitor Camostat. (A) Rhabdoviral pseudotypes bearing the indicated S protein variants, VSV-G or no viral glycoprotein were inoculated onto BHK-21 cells previously transfected with empty plasmid or human angiotensin-converting enzyme 2 (hACE2) expression vector. (B) Rhabdoviral pseudotypes bearing the indicated S protein variants, VSV-G (shown in SI Figure 1) or no viral glycoprotein were inoculated onto 293T, 293T (ACE2), Calu-3, Calu-3 (ACE2), Caco-2, A549-ACE2, Huh-7 (all human) or Vero76 (non-human primate) cells. (C) Rhabdoviral pseudotypes bearing the indicated S protein variants or VSV-G were preincubated with different dilutions of a soluble hACE2 form fused to the Fc portion of human immunoglobulin G (sol-hACE2-Fc) and subsequently inoculated onto Vero76 cells. (D) Rhabdoviral pseudotypes bearing the indicated S protein variants or VSV-G were inoculated onto Calu-3 cells that were preincubated with different concentrations of Camostat. For all panels: Transduction efficiency was quantified at 16 h postinoculation by measuring the activity of virus-encoded luciferase in cell lysates. Presented are the normalized average (mean) data of three biological replicates, each performed with technical quadruplicates. Error bars indicate the standard error of the mean (SEM). Statistical significance was tested by one-(panels a and b) or two-way (panels c and d) ANOVA with Dunnett’s post-hoc test (*P* > 0.05, not significant [ns]; *P* ≤ 0.05, *; *P* ≤ 0.01, **; *P* ≤ 0.001, ***).

We next investigated whether mutations observed in SARS-CoV-2 infected mink altered susceptibility of viral entry to inhibition by soluble ACE2 (Monteil et al., 2020) and Camostat, a protease inhibitor active against TMPRSS2 (Hoffmann et al., 2020). Preincubation of particles bearing S protein with soluble ACE2 and preincubation of Calu-3 lung cells with Camostat efficiently blocked entry driven by all S proteins analyzed (Fig 2C and D), with mutant D614G + cluster 5 being particularly sensitive to inhibition by soluble ACE2 (Fig. 2C). In contrast, entry driven by VSV-G was not affected (Fig 2D-E). Thus, mutations acquired in mink may not compromise SARS-CoV-2 inhibition by Camostat and soluble ACE2.

A high fraction of convalescent COVID-19 patients exhibits a neutralizing antibody response directed against the S protein that may render most of these patients at least temporarily immune to symptomatic reinfection (Rodda et al., 2020; Wajnberg et al., 2020). Similarly, mRNA-based vaccines induce neutralizing antibodies that play an important role in protection from COVID-19 (Polack et al., 2020; Sahin et al., 2020). Finally, neutralizing monoclonal antibodies are currently being developed for COVID-19 therapy and two have received an emergency use authorization (EUA) for COVID-19 therapy (Baum et al., 2020a; Baum et al., 2020b; Hansen et al., 2020). Therefore, we asked whether S protein mutations found in mink compromise SARS-CoV-2 inhibition by serum or plasma from convalescent COVID-19 patients and neutralizing monoclonal antibodies.

We focused our analysis on mutation Y453F, since this mutation is located in the RBD, which constitutes the primary target for neutralizing antibodies. Serum from a control patient failed to inhibit VSV-G or S protein-driven entry (Neg serum #1), as expected. In contrast, 13 out of 14 serum or plasma samples from COVID-19 patients (Pos samples #1-3 and #5-14) potently inhibited S protein but not VSV-G-driven entry while the remaining serum (Pos serum #4) only showed moderate neutralization of S protein-driven entry (Fig. 3A). Importantly, mutation Y453F reduced inhibition by most serum/plasma samples tested, albeit with variable efficiency (median increase of serum/plasma tier required for 50% neutralization [NT50] = 1.62x, range = 1.02x to 3.43x), indicating that this RBD mutation may compromise SARS-CoV-2 control by pre-existing neutralizing antibody responses (Fig. 3A and SI Fig. S2). Similarly, the mutation Y453F reduced inhibition by one (Casirivimab/ REGN10933) out of a cocktail of two antibodies with EUA for COVID-19 therapy (REGN-COV2), while an unrelated, non-neutralizing antibody was inactive (IgG1) (Fig. 3B and SI Fig. S3). Finally, the interference of Y453F with entry inhibition by Casirivimab/REGN10933 was in keeping with position 453 being located at the interface of the S protein and the antibody (SI Fig. S4) and with results reported by a previous study (Baum et al., 2020b). Thus, mutation Y453F that arose in infected mink can compromise viral inhibition by human antibodies induced upon SARS-CoV-2 infection or under development for COVID-19 treatment.

**Figure 3.**
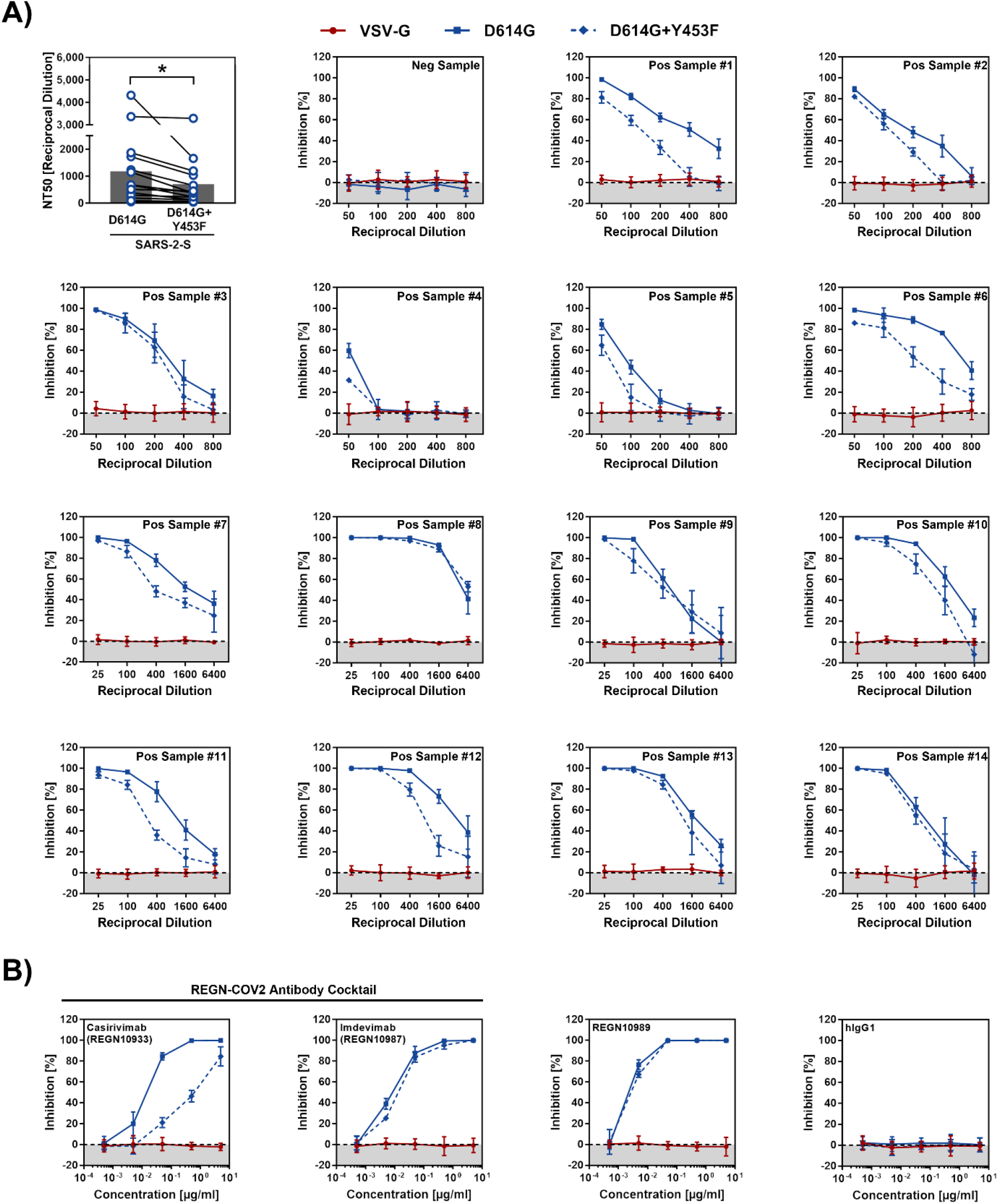
Y453F reduces neutralization by convalescent sera and monoclonal antibodies. (A) Rhabdoviral pseudotypes bearing the indicated spike (S) protein variants or VSV-G were preincubated with different dilutions of serum (Pos Samples #1-6) or plasma (Pos samples #7-14) from convalescent COVID-19 patients (serum from a healthy individual served as control, Neg Sample) before being inoculated onto Vero76 cells. Transduction efficiency was quantified at 16 h postinoculation by measuring the activity of virus-encoded luciferase in cell lysates. The top left panel indicates the serum/plasma titers that lead to a 50% reduction in transduction efficiency (neutralizing titer 50, NT50), which was calculated by a non-linear regression model. Data points from identical serum/plasma samples are connected by lines (grey bars indicate the mean NT50 values for all positive samples). Statistical significance of differences in NT50 values between SARS-2-S harboring D614G alone or in conjunction with Y453F was analyzed by paired student’s t-test (*P* = 0.0212). (B) The experiment outlined in panel A was repeated using serial dilutions of human monoclonal antibodies. For panels A and B: Presented are the normalized average (mean) data of a single experiment performed with technical quadruplicates. Results were confirmed in a separate experiment (due to limited sample material, only two technical replicates could be analyzed in the confirmatory experiment for the serum samples shown in panel A). Error bars indicate the standard deviation.

## DISCUSSION

It is believed that SARS-CoV-2 has been introduced into the human population from an animal reservoir, potentially bats or pangolins (Lam et al., 2020; Xiao et al., 2020; Zhou et al., 2020). Furthermore, the virus can replicate in diverse animal species, including cats, tigers, and minks, for which human-to-animal transmission has been reported (Halfmann et al., 2020; McAloose et al., 2020; Molenaar et al., 2020; Oreshkova et al., 2020; Oude Munnink et al., 2020; Segales et al., 2020; Shi et al., 2020). The virus is likely to acquire adaptive mutations that ensure efficient viral spread in these species, for instance by optimizing interactions with critical host cell factors like the entry receptor ACE2. Indeed, mutation Y453F observed in mink may be an adaptation to efficient use of mink ACE2 for entry, since amino acid 453 is known to make direct contact with human ACE2 (Lan et al., 2020; Wang et al., 2020) and mutation Y453F increases human ACE2 binding (Starr et al., 2020). Moreover, viruses bearing Y453F emerged during experimental infection of ferrets and it has been speculated that Y453F might reflect adaptation of the S protein to ferret ACE2 (Everett et al., 2021). Alternatively, Y453F might be the result of viral evasion of the antibody response and a recent report on emergence of Y453F is a patient with long term COVID-19 supports this possibility (Bazykin, 2021).

The presence of mutation Y453F alone or in combination with H69Δ and V70Δ did not compromise S protein-mediated entry into human cells and its inhibition by soluble ACE2. However, entry into certain cell lines was reduced when Y453F was combined with H69Δ, V70Δ, I692V and M1229I, as found in the S protein of the SARS-CoV-2 cluster 5 variant. This could explain why the cluster 5 variant did not efficiently spread among humans and vanished shortly after its introduction in the human population. The cluster 5 variant S protein was also more sensitive to inhibition by soluble ACE2, hinting towards changes in ACE2 binding affinity when all five signature mutations are present.

Y453F markedly reduced the neutralizing potential of an antibody with an emergency use authorization (Casirivimab/REGN10933). Casirivimab/REGN10933 is one out of two antibodies present in the REGN-COV2 antibody cocktail. The other antibody, Imdevimab/REGN10987, targets a different region in the S protein and inhibited S protein-driven entry with high efficiency regardless of the presence of Y453F. In keeping with this finding, a combination of Casirivimab/REGN10933 and Imdevimab/REGN10987 efficiently blocked SARS-CoV-2 with Y453F in cell culture (Baum et al., 2020b). Maybe more concerning is that Y453F diminished entry inhibition by human sera/plasma from convalescent COVID-19 patients. This finding suggests that at least in a fraction of patients antibody responses induced upon infection and potentially also vaccination might provide only incomplete protection against infection with SARS-CoV-2 amplified in mink. In this context it needs to be stated that most serum/plasma samples analyzed completely inhibited entry at the lowest dilution tested, suggesting that individuals that have high antibody titers (induced upon infection or vaccination) might be protected from infection with mink-derived SARS-CoV-2. The transmission of SARS-CoV-2 to wild minks is another alarming observation (ProMed-mail, 2020h), as such transmission events might generate a permanent natural reservoir for such viruses and new emerging variants that could represent a future threat to wildlife and human health.

The following limitations of our study need to be considered. We employed pseudotyped particles instead of authentic SARS-CoV-2 and we did not determine whether Y453F affects viral inhibition by T cell responses raised against SARS-CoV-2. Further, we did not investigate whether presence of Y453F in the SARS-CoV-2 S protein increases binding to mink ACE2. Nevertheless, our results suggest that the introduction of SARS-CoV-2 into mink allows the virus to acquire mutations that compromise viral control by the humoral immune response in humans. As a consequence, infection of mink and other animal species should be prevented and it should be continuously monitored whether SARS-CoV-2 amplification in other wild or domestic animals occurs and changes critical biological properties of the virus.

## MATERIALS AND METHODS

### Cell culture

All cell lines were incubated at 37 °C in a humidified atmosphere containing 5% CO_2_. 293T (human, kidney; ACC-635, DSMZ), Huh-7 (human, liver; JCRB0403, JCRB; kindly provided by Thomas Pietschmann, TWINCORE, Centre for Experimental and Clinical Infection Research, Hannover, Germany) and Vero76 cells (African green monkey, kidney; CRL-1586, ATCC; kindly provided by Andrea Maisner, Institute of Virology, Philipps University Marburg, Marburg, Germany) were cultivated in Dulbecco’s modified Eagle medium (DMEM) containing 10% fetal bovine serum (FCS, Biochrom), 100 U/ml of penicillin and 0.1 mg/ml of streptomycin (PAN-Biotech). Caco-2 (human, intestine; HTB-37, ATCC) and Calu-3 cells (human, lung; HTB-55, ATCC; kindly provided by Stephan Ludwig, Institute of Virology, University of Münster, Germany) were cultivated in minimum essential medium supplemented with 10% FCS, 100 U/ml of penicillin and 0.1 mg/ml of streptomycin (PAN-Biotech), 1x non-essential amino acid solution (from 100x stock, PAA) and 1 mM sodium pyruvate (Thermo Fisher Scientific). A549 cells (human, lung; CRM-CCL-185, ATCC) were cultivated in DMEM/F-12 medium with Nutrient Mix (Thermo Fisher Scientific) supplemented with 10% FCS, 100 U/ml of penicillin and 0.1 mg/ml of streptomycin (PAN-Biotech). In order to obtain 293T, A549 and Calu-3 cells stably expressing human ACE2, cells were transduced with murine leukemia virus-based transduction vectors and subsequently transduced cells were selected with puromycin (Invivogen). Authentication of cell lines was performed by STR-typing, amplification and sequencing of a fragment of the cytochrome c oxidase gene, microscopic examination and/or according to their growth characteristics. Further, cell lines were routinely tested for contamination by mycoplasma.

### Plasmids

Expression plasmids for vesicular stomatitis virus glycoprotein (VSV-G) (Brinkmann et al., 2017), severe acute respiratory syndrome coronavirus 2 spike glycoprotein (SARS-2-S) containing either a C-terminal HA-epitope tag (SARS-2-S-HA, used for detection in immunoblot) or a truncated cytoplasmic domain (deletion of last 18 amino acid residues at the C-terminus, SARS-2-SΔ18, used for transduction experiments) (Hoffmann et al., 2020) have been described before. Mink-specific mutations were introduced into the expression plasmids for wildtype SARS-2-SΔ18 and SARS-2-S-HA by overlap-extension polymerase chain reaction (PCR) and the resulting PCR products were inserted into the pCG1 expression plasmid (kindly provided by Roberto Cattaneo, Mayo Clinic College of Medicine, Rochester, MN, USA) making use of BamHI and XbaI restriction sites.

In order to obtain the expression plasmid for delivering ACE2 into cell lines via retroviral transduction, the coding sequence for human ACE2 (NM_001371415.1) was inserted into the pQCXIP plasmid (Brass et al., 2009) making use of NotI and PacI restriction sites. Further, we generated an expression plasmid for soluble ACE2 fused to the Fc-portion of human immunoglobulin G (sol-hACE2-Fc). For this, the sequence coding for the ACE2 ectodomain (amino acid residues 1-733) was PCR-amplified and inserted into the pCG1-Fc plasmid (Sauer et al., 2014) (kindly provided by Georg Herrler, University of Veterinary Medicine, Hannover, Germany) making use of PacI and SalI restriction sites. Sequence integrity was verified by sequencing using a commercial sequencing service (Microsynth Seqlab). 293T cells were transfected by calcium-phosphate precipitation, whereas for transfection of BHK-21 cells Lipofectamine LTX with Plus reagent (Thermo Fisher Scientific) was used.

### Sequence analysis and protein models

Spike protein sequences from a total of 742 SARS-CoV-2 isolates were retrieved from the GISAID (global initiative on sharing all influenza data) database (https://www.gisaid.org/) and analyzed regarding the presence of mink-specific mutations. A summary of the selected S protein sequences, including their GISAID accession numbers, is given in SI-Table. Sequence alignments were performed using the Clustal Omega online tool (https://www.ebi.ac.uk/Tools/msa/clustalo/). Protein models were designed using the YASARA (http://www.yasara.org/index.html) and UCSF Chimera (version 1.14, developed by the Resource for Biocomputing, Visualization, and Informatics at the University of California, San Francisco) software packages, and are either based on PDB: 6XDG (Hansen et al., 2020) or on a template generated by modelling the SARS-2-S sequence on a published crystal structure (PDB: 6XR8, (Cai et al., 2020)) with the help of the SWISS-MODEL online tool (https://swissmodel.expasy.org/).

### Patient serum and plasma samples

Serum samples were obtained by the Department of Transfusion Medicine of the University Medical Center Göttingen, Göttingen, Germany. Written consent was obtained from all individuals and the study was approved by the local ethics committee (14/8/20). Collection of plasma samples from COVID-19 patients treated at the intensive care unit was approved by the Ethic committee of the University Medicine Göttingen (SeptImmun Study 25/4/19 Ü). Serum and plasma samples were pre-screened for neutralizing activity using SARS-2-S WT pseudotypes, as described below.

### Production of recombinant human monoclonal antibodies against SARS-CoV-2 spike

VH and VL sequences of Regeneron antibodies Casirivimab/REGN10933, Imdevimab/REGN10987 and REGN10989 (Hansen et al., 2020) were cloned in pCMC3-untagged-NCV (SINO Biologics, Cat: CV011) and produced in 293T cells by SINO Biological (Beijing, China). The human IgG1 isotype control antibodies IgG1/κ and IgG1/λ were produced by transfecting FreeStyle 293-F or 293T cells (Fisher Scientific, Schwerte, Germany, Cat. no. R790-07) with the respective plasmids using the protocol provided with the FreeStyle 293 Expression System (Thermo Fisher Scientific, Cat. no. K9000-01). The isotypes contain human V regions from hybridomas that were established from a human HHKKLL Trianni mouse (Patent US 2013/0219535 A1). Antibodies were affinity-purified from filtered cultured supernatant on a High-Trap protein G column (GE Healthcare, Chicago, USA, Cat.Nr 17-0404-01).

The binding of recombinant antibodies to SARS-2-S was determined by flow cytometry with 293T cells stably transfected with plasmid pWHE469-SARS-CoV2 containing the ORF of the spike protein of SARS-CoV-2 isolate Wuhan-Hu-1 (position 21580 – 25400 from GenBank NC_045512) and a GFP reporter plasmid under the control of a doxycycline-inducible promotor (Krueger et al., 2006). Briefly, 293T cells were stained with the recombinant human IgG1 antibodies in FACS buffer (PBS with 0.5% bovine serum albumin and 1 nmol sodium azide) for 20 minutes in ice, washed, incubated with an Alexa Fluor 647-labeled mouse monoclonal antibody against the human IgG1-Fc (Biolegend, San Diego, USA, cat #409320) and analyzed in a Gallios flow cytometer (Beckman Coulter, Brea, California, USA respectively).

### Production of rhabdoviral pseudotype particles and transduction of target cells

Rhabdoviral pseudotype particles bearing WT or mutant SARS-2-S, VSV-G or no viral protein (negative control) were prepared according to a published protocol (Kleine-Weber et al., 2019) and involved a replication-deficient VSV vector that lacks the genetic information for VSV-G and instead codes for two reporter proteins, enhanced green fluorescent protein and firefly luciferase (FLuc), VSV∗ΔG-FLuc (kindly provided by Gert Zimmer, Institute of Virology and Immunology, Mittelhäusern, Switzerland) (Berger Rentsch and Zimmer, 2011). In brief, 293T cells expressing the desired viral glycoprotein following transfection were inoculated with VSV∗ΔG-FLuc and incubated for 1 h at 37 °C before the inoculum was removed and cells were washed. Finally, culture medium was added that was supplemented with anti-VSV-G antibody (culture supernatant from I1-hybridoma cells; ATCC no. CRL-2700; not added to cells expressing VSV-G). Following an incubation period of 16-18 h, pseudotype particles were harvested by collecting the culture supernatant, pelleting cellular debris through centrifugation (2,000 x g, 10 min, room temperature) and transferring aliquots of the clarified supernatant into fresh reaction tubes. Aliquoted pseudotypes were stored at −80 °C until further use.

For transduction experiments, target cells were seeded into 96-well plates. The following experimental set-ups were used: (i) In case of experiments comparing the efficiency cell entry by WT and mutant SARS-2-S, target cells were inoculated with 100 μl/well of the respective pseudotype particles; (ii) For investigation of inhibition of SARS-2-S-driven cell entry by the serine protease inhibitor Camostat mesylate, Calu-3 cells were preincubated for 1 h with medium (50 μl/well) containing either increasing concentrations of Camostat (0.5, 5 or 50 μM; Tocris) or dimethyl sulfoxide (solvent control) before the respective pseudotype particles were added on top; in order to assess the ability of sol-hACE2-Fc, patient sera and monoclonal antibodies to block SARS-2-S-driven cell entry, pseudotype particles were preincubated for 30 min with medium containing different dilutions of either sol-hACE2-Fc (1:20, 1:200, 1:2,000) or patient serum/plasma (serum: 1:50, 1:100, 1:200, 1:400, 1:800; plasma: 1:25, 1:100, 1:400, 1:1600, 1:6400), or with different concentrations of monoclonal antibody (5, 0.5, 0.05, 0.005, 0.0005 μg/ml), before being inoculated onto Vero76 cells. Pseudotype particles incubated with medium alone served as controls. In all cases, transduction efficiency was analyzed at 16-18 h postinoculation. For this, the culture supernatant was removed and cells were lysed by incubation for 30 min at room temperature with Cell Culture Lysis Reagent (Promega). Next, lysates were transferred into white 96-well plates and FLuc activity was measured using a commercial substrate (Beetle-Juice, PJK) and a Hidex Sense plate luminometer (Hidex).

### Production of sol-hACE2-Fc

293T cells were grown in a T-75 flask and transfected with 20 μg of sol-hACE2-Fc expression plasmid. At 10 h posttransfection, the medium was replaced and cells were further incubated for 38 h before the culture supernatant was collected and centrifuged (2,000 x g, 10 min, 4 °C). Next, the clarified supernatant was loaded onto Vivaspin protein concentrator columns with a molecular weight cut-off of 30 kDa (Sartorius) and centrifuged at 4,000 x g, 4 °C until the sample was concentrated by a factor of 20. The concentrated sol-hACE2-Fc was aliquoted and stored at - 80 ° until further use.

### Analysis of S protein expression, processing and particle incorporation by immunoblot

A total volume of 1 ml of culture medium containing rhabdoviral pseudotypes bearing WT or mutant SARS-2-S-HA were loaded onto a 20% (w/v) sucrose cushion (50 μl) and subjected to high-speed centrifugation (25.000 x g,120 min, 4°C). As controls, particles bearing no S protein or culture medium alone were used. Following centrifugation, 1 ml of supernatant was removed and the residual volume was mixed with 50 μl of 2x SDS-sample buffer (0.03 M Tris-HCl, 10% glycerol, 2% SDS, 0.2% bromophenol blue, 1 mM EDTA) and incubated at 96 °C for 15 min. Next, samples were subjected to SDS-polyacrylamide gel electrophoresis and proteins were blotted onto nitrocellulose membranes using the Mini Trans-Blot Cell system (Bio-Rad).

Following blocking of the membranes by incubation in 5% skim milk solution (skim milk powder dissolved in PBS containing 0.05% Tween-20, PBS-T) for 1 h at room temperature, the membranes were cut in around the 55 kDa marker band of the protein marker (PageRuler Prestained Protein Ladder, Thermo Fisher Scientific). The upper portion of the membrane was probed with anti-HA tag antibody (mouse, Sigma-Aldrich, H3663) diluted 1:1,000 in 5% skim milk solution, while the lower portion of the membrane was probed with anti-VSV matrix protein antibody (Kerafast, EB0011; loading control) diluted 1:2,500 in 5% skim milk solution. Following incubation over night at 4 °C, membranes were washed three times with PBS-T, before being probed with peroxidase-conjugated anti-mouse antibody (Dianova, 115-035-003, 1:5,000) for 1 h at room temperature. Thereafter, the membranes were washed again three times with PBS-T, incubated with an in house-prepared developing solution (1 ml of solution A: 0.1 M Tris-HCl [pH 8.6], 250 μg/ml luminol sodium salt; 100 μl of solution B: 1 mg/ml para-hydroxycoumaric acid dissolved in dimethyl sulfoxide [DMSO]; 1.5 μl of 0.3 % H_2_O_2_ solution) and imaged using the ChemoCam imager along with the ChemoStar Imager Software version v.0.3.23 (Intas Science Imaging Instruments GmbH).

### Data normalization and statistical analysis

Data analysis was performed using Microsoft Excel as part of the Microsoft Office software package (version 2019, Microsoft Corporation) and GraphPad Prism 8 version 8.4.3 (GraphPad Software). Data normalization was done as follows: (i) In order to assess enhancement of S protein-driven pseudotype entry in BHK-21 cells following directed overexpression of hACE2, transduction was normalized against the assay background (which was determined by using rhabdoviral pseudotypes bearing no viral glycoprotein, set as 1); (ii) To compare efficiency of cell entry driven by the different S protein variants under study, transduction was normalized against SARS-2-S WT (set as 100%); (iii) For experiments investigating inhibitory effects exerted by sol-hACE2-Fc or Camostat Mesylate, patient serum/plasma samples or monoclonal antibodies, transduction was normalized against a reference sample (control-treated cells or pseudotypes, set as 100%). Statistical significance was tested by one- or two-way analysis of variance (ANOVA) with Dunnett’s or Sidak’s post-hoc test or by paired student’s t-test. Only *P* values of 0.05 or lower were considered statistically significant (*P* > 0.05, not significant [ns]; *P* ≤ 0.05, *; *P* ≤ 0.01, **; *P* ≤ 0.001, ***). Specific details on the statistical test and the error bars are indicated in the figure legends. NT50 (neutralizing titer 50) values, which indicate the serum/plasma titers that lead to a 50% reduction in transduction efficiency, were calculated using a non-linear regression model.

## SUPPLEMENTAL INFORMATION

**SI Table.** Summary of S protein sequences used for analysis and their respective sequence information (related to Figure 1C).

**Figure S1.**
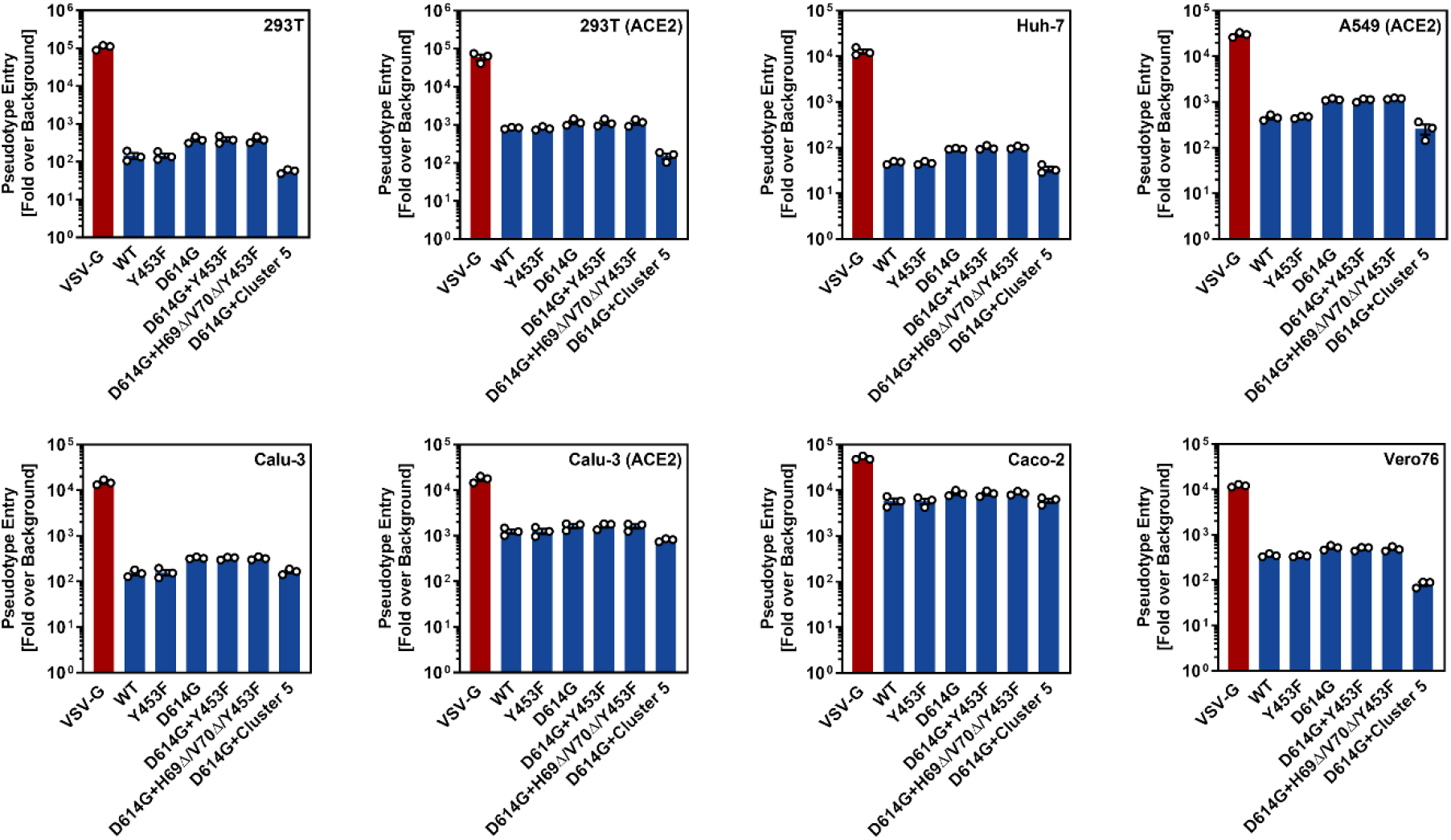
Transduction of target cells (related to Figure 2B). Data presented in Figure 2B were normalized against the assay background (set as 1). Further, transduction efficiency by pseudotype particles bearing VSV-G is shown.

**Figure S2.**
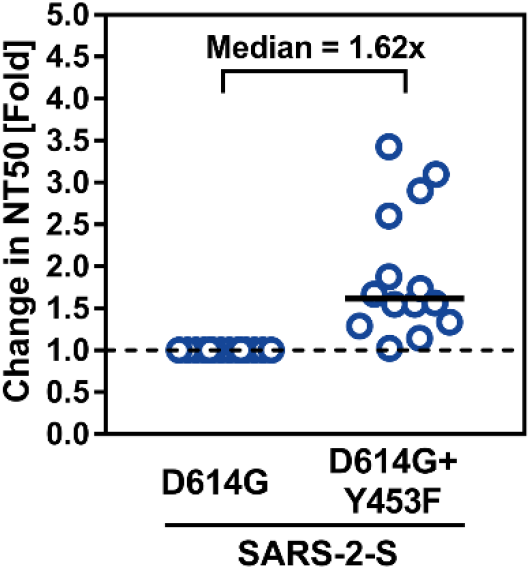
Presence of Y453T reduces antibody-mediated neutralization (related to Figure 3A). The relative difference in NT50 values between SARS-2-S harboring D614G alone or in conjunction with Y453F was calculated (indicated as Fold difference with SARS-2-S D614G set as 1). The median is indicated by a black line.

**Figure S3.**
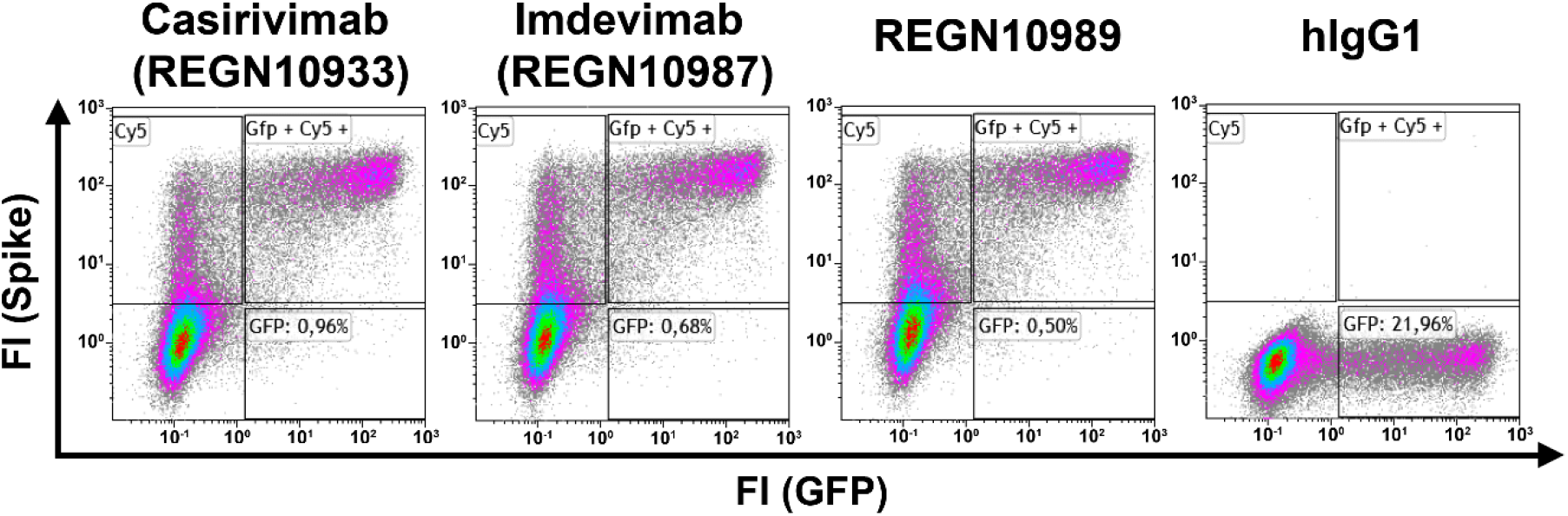
Flow cytometric detection of antibody-binding to cell-expressed SARS-2-S (related to Figure 3B). 293T cells stably transfected with a doxycycline-inducible SARS-2-S (hCoV-19/Wuhan/Hu-1/2019 hCoV-19/Wuhan/Hu-1/2019 isolate) were stained with the indicated Regeneron (REGN) antibodies and an Alexa Fluor 643-conjugated anti-human IgG antibody. A recombinant human IgG served as an isotype control. FI, fluorescence intensity.

**Figure S4.**
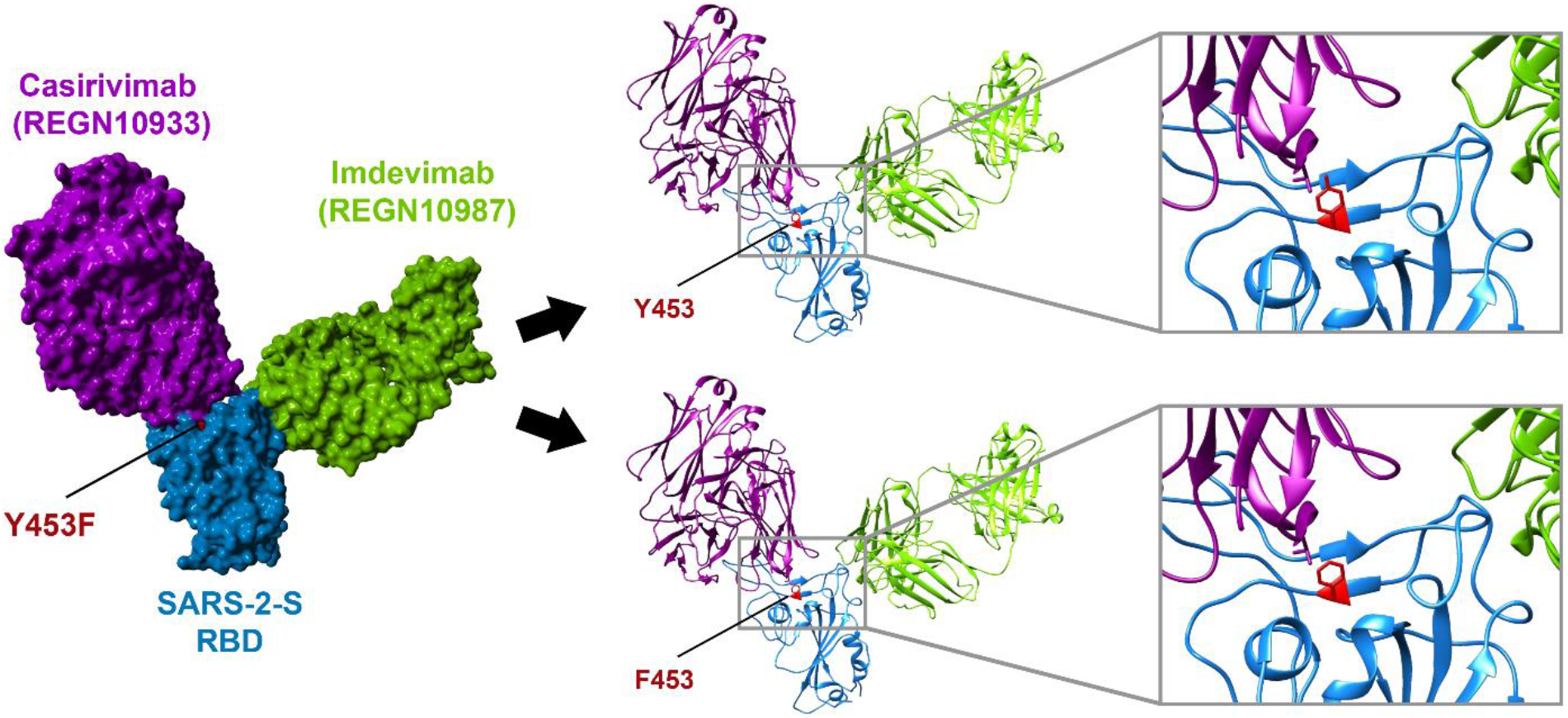
Y453F centers in the binding interface of antibody REGN10933 and the SARS-2-S RBD (related to Figure 3B). The protein models of the SARS-2-S receptor-binding domain (RBD, blue) in complex with antibodies REGN10933 (purple) and REGN10987 (green) were constructed based on the 6XDG template (Hansen et al., 2020). Residues highlighted in red indicate amino acid position 453 in SARS-2-S RBD (either tyrosine [Y] or phenylalanine [F]).

## ACKNOWLEDGMENTS

We like to thank Roberto Cattaneo, Georg Herrler, Stephan Ludwig, Andrea Maisner, Thomas Pietschmann and Gert Zimmer for providing reagents and Tobit Steinmetz for the 293T cell line that stably expresses SARS-2-S. Further, we thank Anna-Sophie Moldenhauer, Sara Krause and Manuela Hauke for excellent technical support. We gratefully acknowledge the originating laboratories responsible for obtaining the specimens and the submitting laboratories where genetic sequence data were generated and shared via the GISAID Initiative, on which this research is based. The Pöhlmann lab is supported by BMBF (RAPID Consortium, 01KI1723D and 01KI2006D; RENACO, 01KI20328A, 01KI20396, COVIM consortium). The Jäck lab is supported by grants from the DFG (grants GRK1660 and TRR130), BMBF (COVIM), and the Bavarian Ministry of Science and Art. The Nessler lab is supported by BMBF (Netzwerk Universitätsmedizin, NUM).

## AUTHOR CONTRIBUTIONS

Conceptualization, M.H., H.-M.J., S.P.; Funding acquisition, S.P.; Investigation, M.H., L.Z., N.K., L.G., H.K.-W., S.S.; Essential resources, H.H.-W., A.K., M.S.W., S.N., J.R., H.-M.J..; Writing, M.H. and S.P., Review and editing, all authors.

## REFERENCES

(WHO), W.H.O. (2021). Weekly operational update on COVID-19 - 9 February 2020 (WHO).

Baum, A., Ajithdoss, D., Copin, R., Zhou, A., Lanza, K., Negron, N., Ni, M., Wei, Y., Mohammadi, K., Musser, B., et al. (2020a). REGN-COV2 antibodies prevent and treat SARS-CoV-2 infection in rhesus macaques and hamsters. Science 370, 1110–1115.

Baum, A., Fulton, B.O., Wloga, E., Copin, R., Pascal, K.E., Russo, V., Giordano, S., Lanza, K., Negron, N., Ni, M., et al. (2020b). Antibody cocktail to SARS-CoV-2 spike protein prevents rapid mutational escape seen with individual antibodies. Science 369, 1014–1018.

Bazykin, G.A.S., O.; Danilenko, D; Fadeev, A; Komissarova, K; Ivanova, A; Sergeeva, M.; Safina, K; Nabieva, E.; Klink, G.; Garushyants, S.; Zabutova, J.; Kholodnaia, A.; Skorokhod, I.; Ryabchikova, V.V.; Komissarov, A.; Lioznov, D. (2021). Emergence of Y453F and Δ69-70HV mutations in a lymphoma patient with long-term COVID-19

Berger Rentsch, M., and Zimmer, G. (2011). A vesicular stomatitis virus replicon-based bioassay for the rapid and sensitive determination of multi-species type I interferon. PLoS One 6, e25858.

Brass, A.L., Huang, I.C., Benita, Y., John, S.P., Krishnan, M.N., Feeley, E.M., Ryan, B.J., Weyer, J.L., van der Weyden, L., Fikrig, E., et al. (2009). The IFITM proteins mediate cellular resistance to influenza A H1N1 virus, West Nile virus, and dengue virus. Cell 139, 1243–1254.

Brinkmann, C., Hoffmann, M., Lubke, A., Nehlmeier, I., Kramer-Kuhl, A., Winkler, M., and Pohlmann, S. (2017). The glycoprotein of vesicular stomatitis virus promotes release of virus-like particles from tetherin-positive cells. PLoS One 12, e0189073.

Cai, Y., Zhang, J., Xiao, T., Peng, H., Sterling, S.M., Walsh, R.M., Jr., Rawson, S., Rits-Volloch, S., and Chen, B. (2020). Distinct conformational states of SARS-CoV-2 spike protein. Science 369, 1586–1592.

Everett, H.E., Lean, F.Z.X., Byrne, A.M.P., van Diemen, P.M., Rhodes, S., James, J., Mollett, B., Coward, V.J., Skinner, P., Warren, C.J., et al. (2021). Intranasal Infection of Ferrets with SARS-CoV-2 as a Model for Asymptomatic Human Infection. Viruses 13.

Guan, Y., Zheng, B.J., He, Y.Q., Liu, X.L., Zhuang, Z.X., Cheung, C.L., Luo, S.W., Li, P.H., Zhang, L.J., Guan, Y.J., et al. (2003). Isolation and characterization of viruses related to the SARS coronavirus from animals in southern China. Science 302, 276–278.

Halfmann, P.J., Hatta, M., Chiba, S., Maemura, T., Fan, S., Takeda, M., Kinoshita, N., Hattori, S.I., Sakai-Tagawa, Y., Iwatsuki-Horimoto, K., et al. (2020). Transmission of SARS-CoV-2 in Domestic Cats. N Engl J Med 383, 592–594.

Hansen, J., Baum, A., Pascal, K.E., Russo, V., Giordano, S., Wloga, E., Fulton, B.O., Yan, Y., Koon, K., Patel, K., et al. (2020). Studies in humanized mice and convalescent humans yield a SARS-CoV-2 antibody cocktail. Science 369, 1010–1014.

Hoffmann, M., Kleine-Weber, H., Schroeder, S., Kruger, N., Herrler, T., Erichsen, S., Schiergens, T.S., Herrler, G., Wu, N.H., Nitsche, A., et al. (2020). SARS-CoV-2 Cell Entry Depends on ACE2 and TMPRSS2 and Is Blocked by a Clinically Proven Protease Inhibitor. Cell 181, 271–280 e278.

Kleine-Weber, H., Elzayat, M.T., Wang, L., Graham, B.S., Muller, M.A., Drosten, C., Pohlmann, S., and Hoffmann, M. (2019). Mutations in the Spike Protein of Middle East Respiratory Syndrome Coronavirus Transmitted in Korea Increase Resistance to Antibody-Mediated Neutralization. J Virol 93.

Koopmans, M. (2020). SARS-CoV-2 and the human-animal interface: outbreaks on mink farms. Lancet Infect Dis.

Korber, B., Fischer, W.M., Gnanakaran, S., Yoon, H., Theiler, J., Abfalterer, W., Hengartner, N., Giorgi, E.E., Bhattacharya, T., Foley, B., et al. (2020). Tracking Changes in SARS-CoV-2 Spike: Evidence that D614G Increases Infectivity of the COVID-19 Virus. Cell 182, 812–827 e819.

Krueger, C., Danke, C., Pfleiderer, K., Schuh, W., Jack, H.M., Lochner, S., Gmeiner, P., Hillen, W., and Berens, C. (2006). A gene regulation system with four distinct expression levels. J Gene Med 8, 1037–1047.

Lam, T.T., Jia, N., Zhang, Y.W., Shum, M.H., Jiang, J.F., Zhu, H.C., Tong, Y.G., Shi, Y.X., Ni, X.B., Liao, Y.S., et al. (2020). Identifying SARS-CoV-2-related coronaviruses in Malayan pangolins. Nature 583, 282–285.

Lan, J., Ge, J., Yu, J., Shan, S., Zhou, H., Fan, S., Zhang, Q., Shi, X., Wang, Q., Zhang, L., et al. (2020). Structure of the SARS-CoV-2 spike receptor-binding domain bound to the ACE2 receptor. Nature 581, 215–220.

Lau, S.K., Woo, P.C., Li, K.S., Huang, Y., Tsoi, H.W., Wong, B.H., Wong, S.S., Leung, S.Y., Chan, K.H., and Yuen, K.Y. (2005). Severe acute respiratory syndrome coronavirus-like virus in Chinese horseshoe bats. Proc Natl Acad Sci U S A 102, 14040–14045.

Leste-Lasserre, C. (2020). Pandemic dooms Danish mink-and mink research. Science 370, 754.

Li, W., Shi, Z., Yu, M., Ren, W., Smith, C., Epstein, J.H., Wang, H., Crameri, G., Hu, Z., Zhang, H., et al. (2005). Bats are natural reservoirs of SARS-like coronaviruses. Science 310, 676–679.

McAloose, D., Laverack, M., Wang, L., Killian, M.L., Caserta, L.C., Yuan, F., Mitchell, P.K., Queen, K., Mauldin, M.R., Cronk, B.D., et al. (2020). From People to Panthera: Natural SARS-CoV-2 Infection in Tigers and Lions at the Bronx Zoo. mBio 11.

Molenaar, R.J., Vreman, S., Hakze-van der Honing, R.W., Zwart, R., de Rond, J., Weesendorp, E., Smit, L.A.M., Koopmans, M., Bouwstra, R., Stegeman, A., et al. (2020). Clinical and Pathological Findings in SARS-CoV-2 Disease Outbreaks in Farmed Mink (Neovison vison). Vet Pathol 57, 653–657.

Monteil, V., Kwon, H., Prado, P., Hagelkruys, A., Wimmer, R.A., Stahl, M., Leopoldi, A., Garreta, E., Hurtado Del Pozo, C., Prosper, F., et al. (2020). Inhibition of SARS-CoV-2 Infections in Engineered Human Tissues Using Clinical-Grade Soluble Human ACE2. Cell 181, 905–913 e907.

Oreshkova, N., Molenaar, R.J., Vreman, S., Harders, F., Oude Munnink, B.B., Hakze-van der Honing, R.W., Gerhards, N., Tolsma, P., Bouwstra, R., Sikkema, R.S., et al. (2020). SARS-CoV-2 infection in farmed minks, the Netherlands, April and May 2020. Euro Surveill 25.

Oude Munnink, B.B., Sikkema, R.S., Nieuwenhuijse, D.F., Molenaar, R.J., Munger, E., Molenkamp, R., van der Spek, A., Tolsma, P., Rietveld, A., Brouwer, M., et al. (2020). Transmission of SARS-CoV-2 on mink farms between humans and mink and back to humans. Science.

Plante, J.A., Liu, Y., Liu, J., Xia, H., Johnson, B.A., Lokugamage, K.G., Zhang, X., Muruato, A.E., Zou, J., Fontes-Garfias, C.R., et al. (2020). Spike mutation D614G alters SARS-CoV-2 fitness. Nature.

Polack, F.P., Thomas, S.J., Kitchin, N., Absalon, J., Gurtman, A., Lockhart, S., Perez, J.L., Perez Marc, G., Moreira, E.D., Zerbini, C., et al. (2020). Safety and Efficacy of the BNT162b2 mRNA Covid-19 Vaccine. N Engl J Med.

ProMed-mail (2020a). CORONAVIRUS DISEASE 2019 UPDATE (135): NETHERLANDS (NORTH BRABANT) ANIMAL, FARMED MINK.

ProMed-mail (2020b). CORONAVIRUS DISEASE 2019 UPDATE (266): DENMARK (NORTH JUTLAND) ANIMAL, FARMED MINK, FIRST REPORT.

ProMed-mail (2020c). CORONAVIRUS DISEASE 2019 UPDATE (366): ANIMAL, USA (UTAH) MINK.

ProMed-mail (2020d). CORONAVIRUS DISEASE 2019 UPDATE (468): ANIMAL, SWEDEN, MINK, FIRST REPORT, OIE.

ProMed-mail (2020e). CORONAVIRUS DISEASE 2019 UPDATE (490): ANIMAL, GREECE (EM) MINK, FIRST REPORT, OIE, ASSESSMENT.

ProMed-mail (2020f). CORONAVIRUS DISEASE 2019 UPDATE (510): ANIMAL, MINK, LITHUANIA, POLAND, FIRST REPORTS, FRANCE, OIE.

ProMed-mail (2020g). CORONAVIRUS DISEASE 2019 UPDATE (531): ANIMAL, CANADA (BRITISH COLUMBIA) MINK, OIE.

ProMed-mail (2020h). CORONAVIRUS DISEASE 2019 UPDATE (536): ANIMAL, USA (UTAH) WILD MINK, FIRST CASE.

Rodda, L.B., Netland, J., Shehata, L., Pruner, K.B., Morawski, P.A., Thouvenel, C.D., Takehara, K.K., Eggenberger, J., Hemann, E.A., Waterman, H.R., et al. (2020). Functional SARS-CoV-2-Specific Immune Memory Persists after Mild COVID-19. Cell.

Sahin, U., Muik, A., Vogler, I., Derhovanessian, E., Kranz, L.M., Vormehr, M., Quandt, J., Bidmon, N., Ulges, A., Baum, A., et al. (2020). BNT162b2 induces SARS-CoV-2-neutralising antibodies and T cells in humans. medRxiv, 2020.2012.2009.20245175.

Sauer, A.K., Liang, C.H., Stech, J., Peeters, B., Quere, P., Schwegmann-Wessels, C., Wu, C.Y., Wong, C.H., and Herrler, G. (2014). Characterization of the sialic acid binding activity of influenza A viruses using soluble variants of the H7 and H9 hemagglutinins. PLoS One 9, e89529.

Segales, J., Puig, M., Rodon, J., Avila-Nieto, C., Carrillo, J., Cantero, G., Terron, M.T., Cruz, S., Parera, M., Noguera-Julian, M., et al. (2020). Detection of SARS-CoV-2 in a cat owned by a COVID-19-affected patient in Spain. Proc Natl Acad Sci U S A 117, 24790–24793.

Shi, J., Wen, Z., Zhong, G., Yang, H., Wang, C., Huang, B., Liu, R., He, X., Shuai, L., Sun, Z., et al. (2020). Susceptibility of ferrets, cats, dogs, and other domesticated animals to SARS-coronavirus 2. Science 368, 1016–1020.

Starr, T.N., Greaney, A.J., Hilton, S.K., Ellis, D., Crawford, K.H.D., Dingens, A.S., Navarro, M.J., Bowen, J.E., Tortorici, M.A., Walls, A.C., et al. (2020). Deep Mutational Scanning of SARS-CoV-2 Receptor Binding Domain Reveals Constraints on Folding and ACE2 Binding. Cell 182, 1295–1310 e1220.

Wajnberg, A., Amanat, F., Firpo, A., Altman, D.R., Bailey, M.J., Mansour, M., McMahon, M., Meade, P., Mendu, D.R., Muellers, K., et al. (2020). Robust neutralizing antibodies to SARS-CoV-2 infection persist for months. Science 370, 1227–1230.

Wang, Q., Zhang, Y., Wu, L., Niu, S., Song, C., Zhang, Z., Lu, G., Qiao, C., Hu, Y., Yuen, K.Y., et al. (2020). Structural and Functional Basis of SARS-CoV-2 Entry by Using Human ACE2. Cell 181, 894–904 e899.

Xiao, K., Zhai, J., Feng, Y., Zhou, N., Zhang, X., Zou, J.J., Li, N., Guo, Y., Li, X., Shen, X., et al. (2020). Isolation of SARS-CoV-2-related coronavirus from Malayan pangolins. Nature 583, 286–289.

Zhou, P., Yang, X.L., Wang, X.G., Hu, B., Zhang, L., Zhang, W., Si, H.R., Zhu, Y., Li, B., Huang, C.L., et al. (2020). A pneumonia outbreak associated with a new coronavirus of probable bat origin. Nature 579, 270–273.

